## Supplementary Figures 1 - 11 and the figure legends Supplementary Table 1 for "Voltage Sensor Conformations Induced by LQTS-associated Mutations in hERG Potassium Channels"

### Supplementary Information for "Voltage Sensor Conformations Induced by LQTS-associated Mutations in hERG Potassium Channels"

Aaron N. Chan[2], Co D. Quach[2], Lucas J. Handlin[1], Erin N. Lessie[1], Emad Tajkhorshid[2], and Gucan Dai[1,*]

Author Affiliations:

[1]Department of Biochemistry and Molecular Biology, Saint Louis University School of Medicine

[2]Theoretical and Computational Biophysics Group, NIH Center for Macromolecular Modeling and Bioinformatics, Beckman Institute for Advanced Science and Technology, Department of Biochemistry, Center for Biophysics and Quantitative Biology, University of Illinois at Urbana-Champaign

This file includes:
Supplementary Figures 1 - 11 and the figure legends
Supplementary Table 1

**Figure S1**

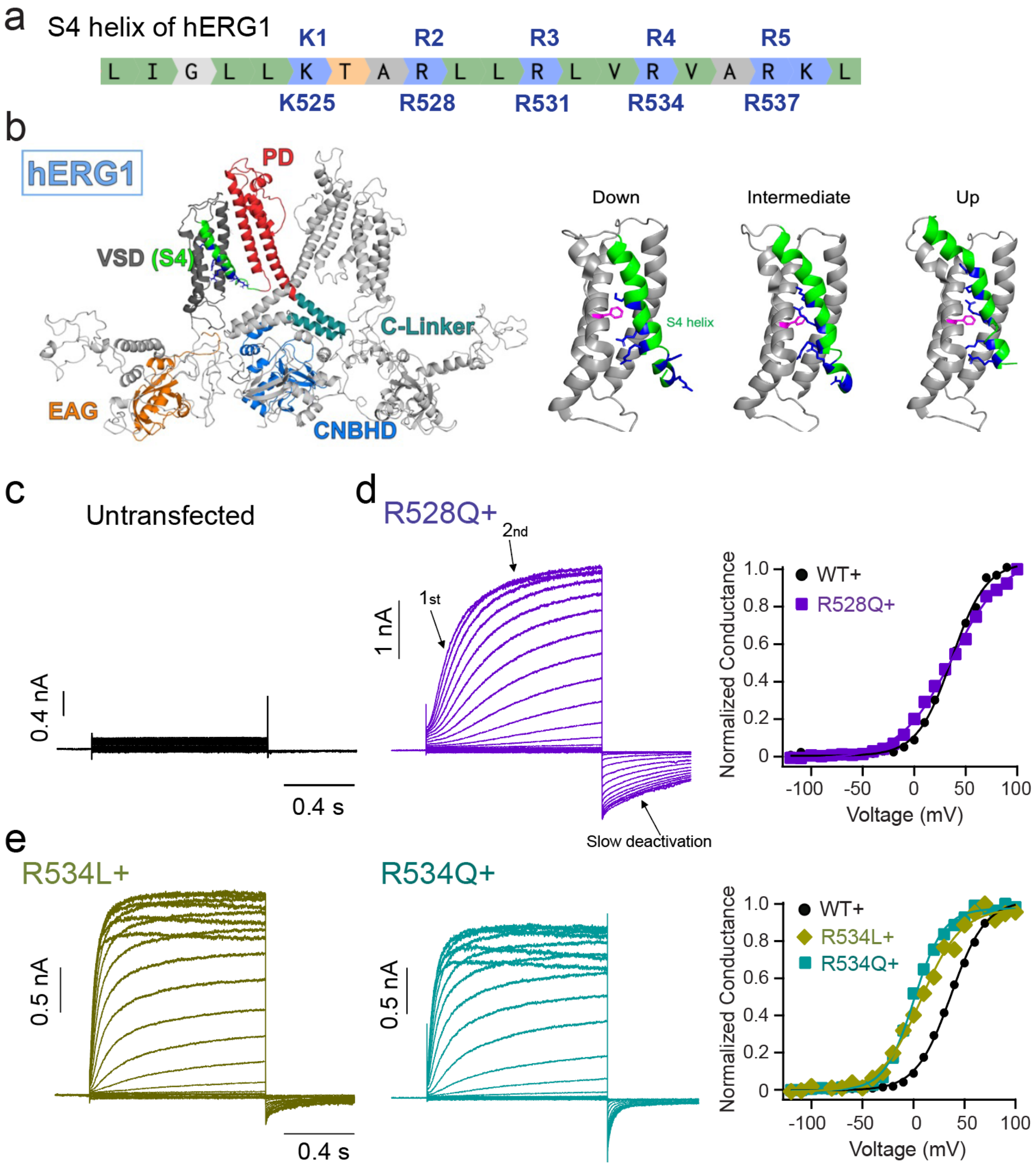


**Supplementary Fig. 1. Activation of hERG channels with mutations of voltage-sensing arginines in the S4 helix.**

**a**. Sequence of the S4 helix in hERG1 channels, with voltage-sensing arginines highlighted. The positions K1 to R5 follow the classic nomenclature based on their analogs in the Shaker potassium channel.

**b**. Left: Architecture of hERG channels, only two subunits of the tetramer are shown. Right: Homology models of the voltage-sensing domain (VSD, S1-S4 helices) of hERG channels in the down, intermediate and up conformations, based on cryo-EM structures of rat Eag channels (PDB codes: 8EP1, 8EP0, and 8EOW respectively). Other parts of the channel are not shown. Positively charged residues of the S4 helix are in blue. The F463 in the charge transfer center is in magenta.
**c.** Representative current traces of patch-clamp recordings of un-transfected tsA cells in a whole-cell configuration, in the presence of 10 mM TEA.
**d-e.** Representative current traces and G-V relationships of hERG-R534L, hERG-R534Q, and hERG-R528Q were shown. Same voltage protocol as in the Fig. 1a was used. For the R528Q mutation, the initial phase made the activation sigmoidal, and the main phase had two components. Note all hERG1 channels in this paper contain the S620T mutation to abolish the fast inactivation.

**Figure S2**

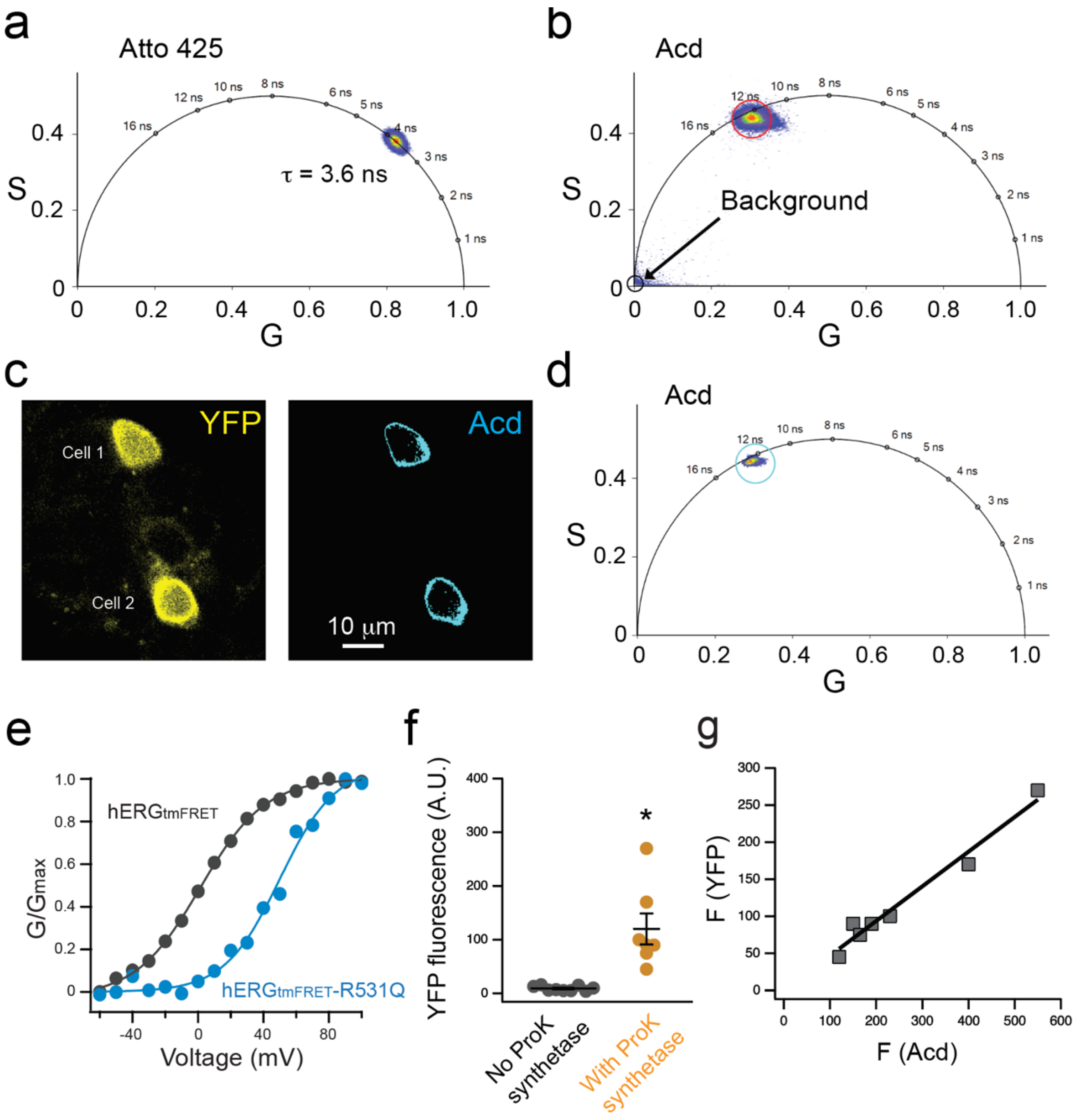


**Supplementary Fig. 2. Illustrations of FLIM calibration, background lifetime species and correlation of Acd with YFP.**

**a.** Phasor plot depicting the lifetime of the calibration dye Atto 425 in water, with a known lifetime of 3.6 ns.

**b.** Phasor plot displaying the background lifetime species associated with the Acd donor alone lifetime species, positioned near the (0,0) coordinate, and before the intensity-based threshold filtering. It is measured from a single tsA cell in the absence of FRET acceptors.

**c.** Left: Representative intensity-based imaging showing YFP fluorescence from two tsA cells in a cluster of cells, indicating the expression of full-length hERG channels. Right: Lifetime-based imaging showing Acd fluorescence at the membrane post-intensity-based selection in reference to the YFP fluorescence localization, eliminating cytosolic Acd fluorescence.

**d.** Phasor plot displaying the combined lifetime species originating from membrane-localized Acd in the same two cells depicted in panel c, after removing the background and cytosolic fluorescence using the intensity-based filtering.
**e**. G-V relationships of $hERGt_{mFRET}$ and $hERG_{tmFRET}$-R531Q channels recorded from tsA cells.
**f**. comparison of membrane localized YFP fluorescence intensity comparing cells with (n = 7 cells) or without (n = 9 cells) transfecting the plasmid encoding the tRNA (M15-PylT)/PylRS pair for incorporating ProK. Data shown are presented as mean ± s.e.m., *$p < 0.05$.
**g**. Linear correlation between the intensities of membrane localized YFP versus Acd fluorescence from individual cells, indicating that fluorescence of Acd originates from the Acd incorporated into YFP-tagged hERG channels.

**Figure S3**

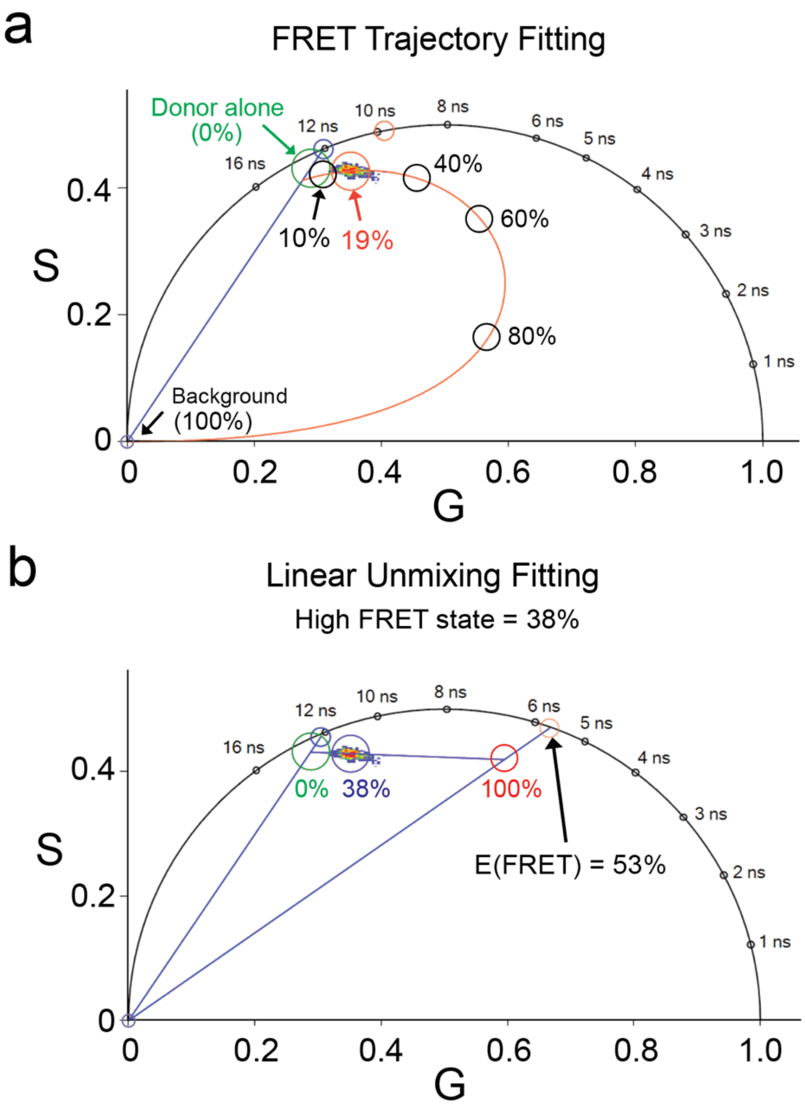


**Supplementary Fig. 3. Comparison of linear unmixing and FRET trajectory methods for phasor FLIM analysis using the VistaVision software.**

**a**. FRET trajectory analysis uses only the center position of the same measured phasor shown in panel a, and in this example, assuming that all donors undergo FRET with acceptors. The red trajectory curve begins at 0% FRET efficiency (donor only) and progresses through 10%, 19% (measured phasor), 40%, 60%, 80% and ends at 100% efficiency. The background contribution was modeled at the (0,0) coordinate, representing ambient light noise, which overlaps the 100% FRET efficiency phasor. This method estimates an average FRET efficiency across the ensemble but does not resolve the fractional contributions of distinct FRET states. As a result, the apparent FRET efficiency is underestimated compared to that obtained in panel b.

**b**. Linear unmixing analysis assuming two donor lifetime populations: a low FRET state with negligible energy transfer, and a high FRET state with a FRET efficiency of 53%. This method estimates the fraction of donors in the high FRET state—38% in this example—by leveraging the spatial distribution pattern of the lifetime species in the plot. A higher proportion of unquenched donor will lead to greater underestimation of the high FRET state fraction.

**Figure S4**

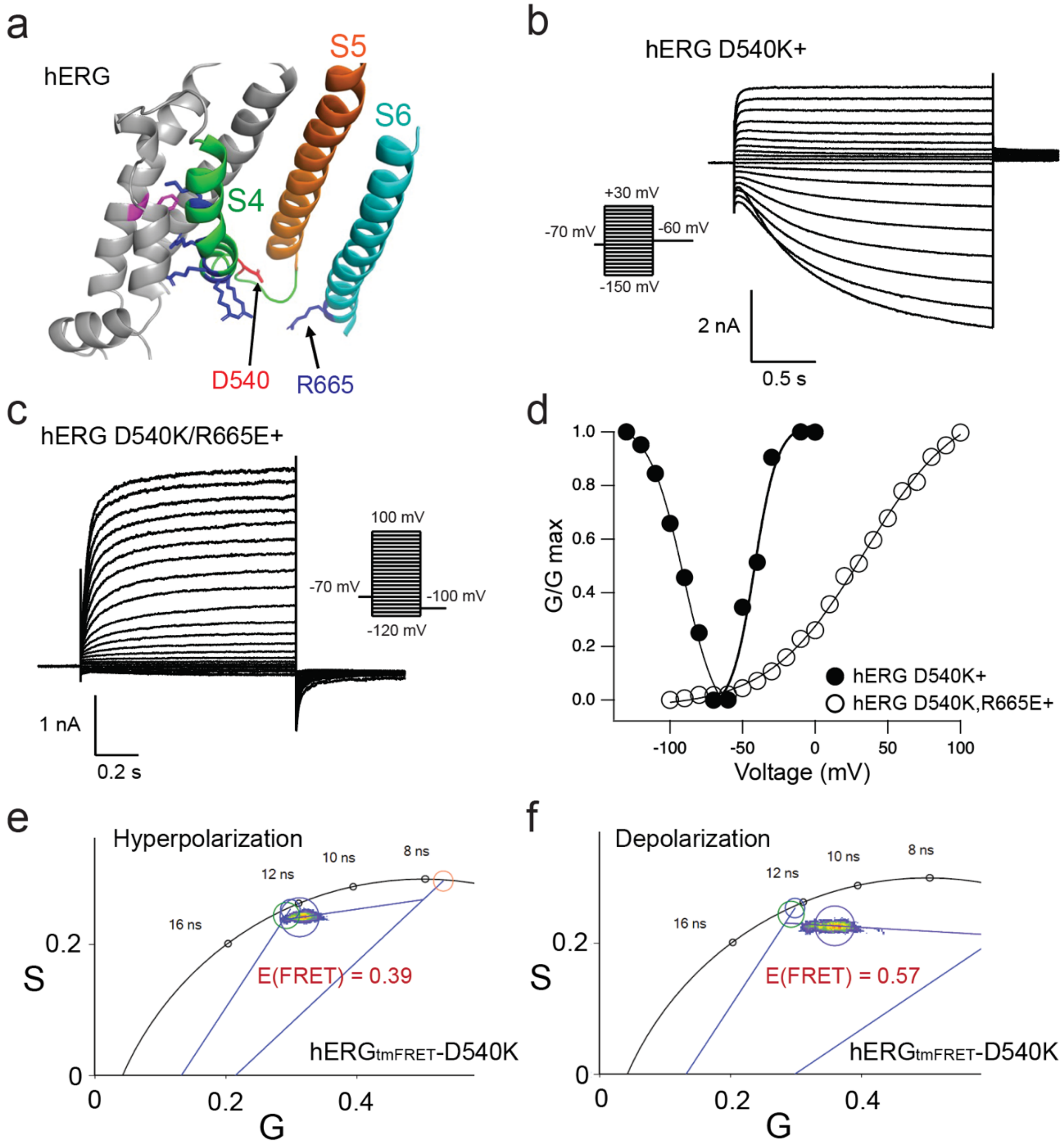


**Supplementary Fig. 4. FLIM-tmFRET measurement of the voltage sensor S4 rearrangement in hERG channels: effects of disrupting the D540/R665 salt bridge.**

**a**. Structural cartoon (upward view) showing the positions of D540 and R665 within the hERG channel.

**b-c**. Representative current traces for activation of hERG-D540K and hERG-D540K/R665E charge-swapped channels.

**d**. Comparison of G-V relationships for activation of hERG-D540K and hERG-D540K/R665E channels.

**e-f**. Combined fluorescence signals from four cells for linear unmixing fitting of hERG-D540K channels in either hyperpolarizing or depolarizing voltage conditions. Hyperpolarization was achieved by extracellular 107 mM NMDG, while depolarization was achieved by extracellular 120 mM KCl. Different FRET efficiencies exhibited when comparing the results from the hyperpolarizing and depolarizing conditions.

**Figure S5**

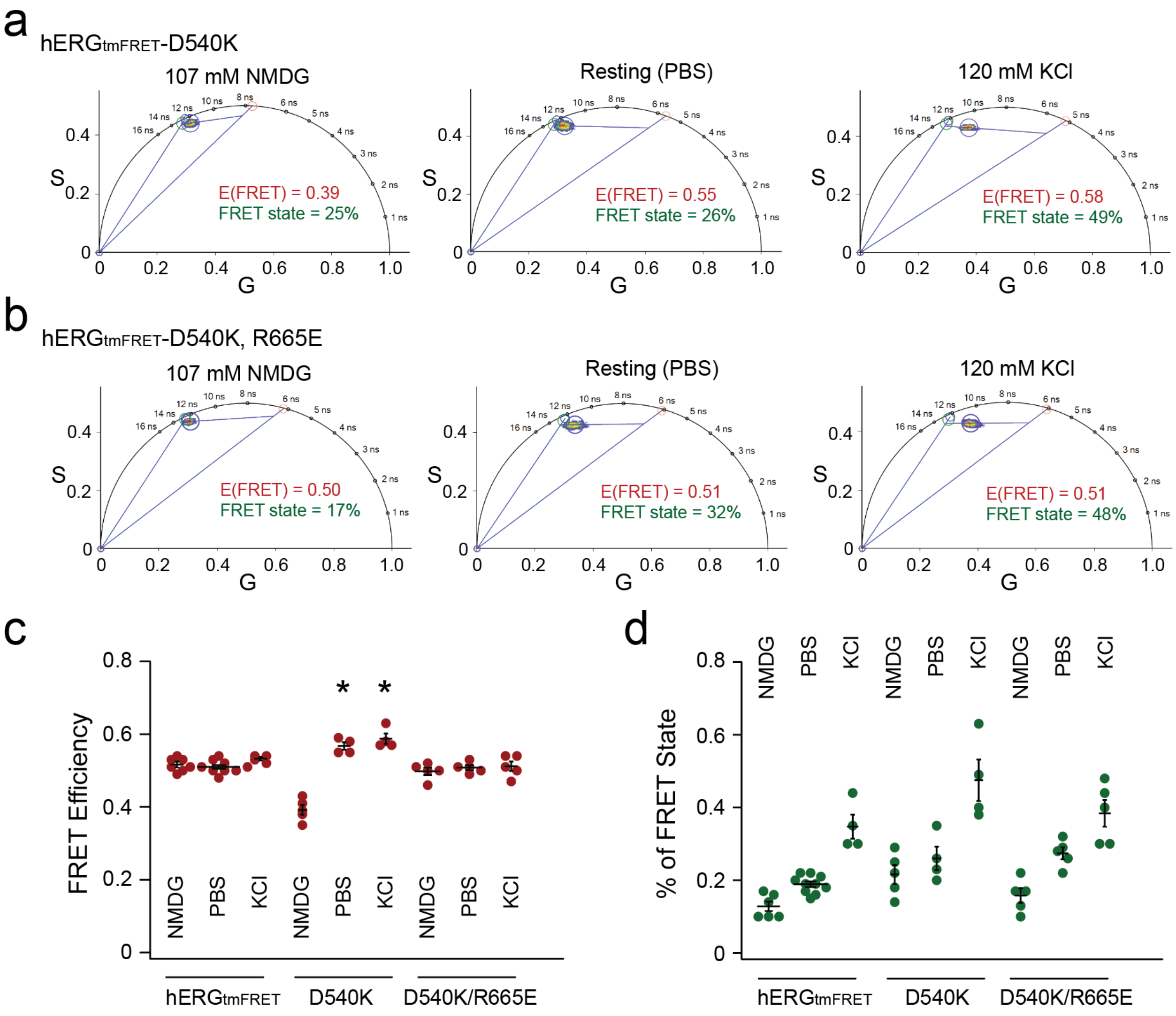


**Supplementary Fig. 5. Phasor linear unmixing analysis of FLIM-tmFRET in hERG channels, after disruption and rescue of the D540/R665 salt bridge.**

**a-b**. Representative analysis of the L523/G439 tmFRET from individual cells for $hERG_{tmFRET}$, $hERG_{tmFRET}$-D540K, and $hERG_{tmFRET}$-D540K/R665E channels under hyperpolarization, resting and depolarization conditions.

**c-d**. Summary data of the estimated FRET efficiencies and percentage of the FRET state using linear unmixing of the phasor approach. Data shown are presented as mean ± s.e.m., n = 4 – 10. For FRET efficiencies of D540K, p = 6e-6 between NMDG and PBS conditions, p = 2e-6 between NMDG and KCl conditions, one-way ANOVA (no adjustment).

**Figure S6**

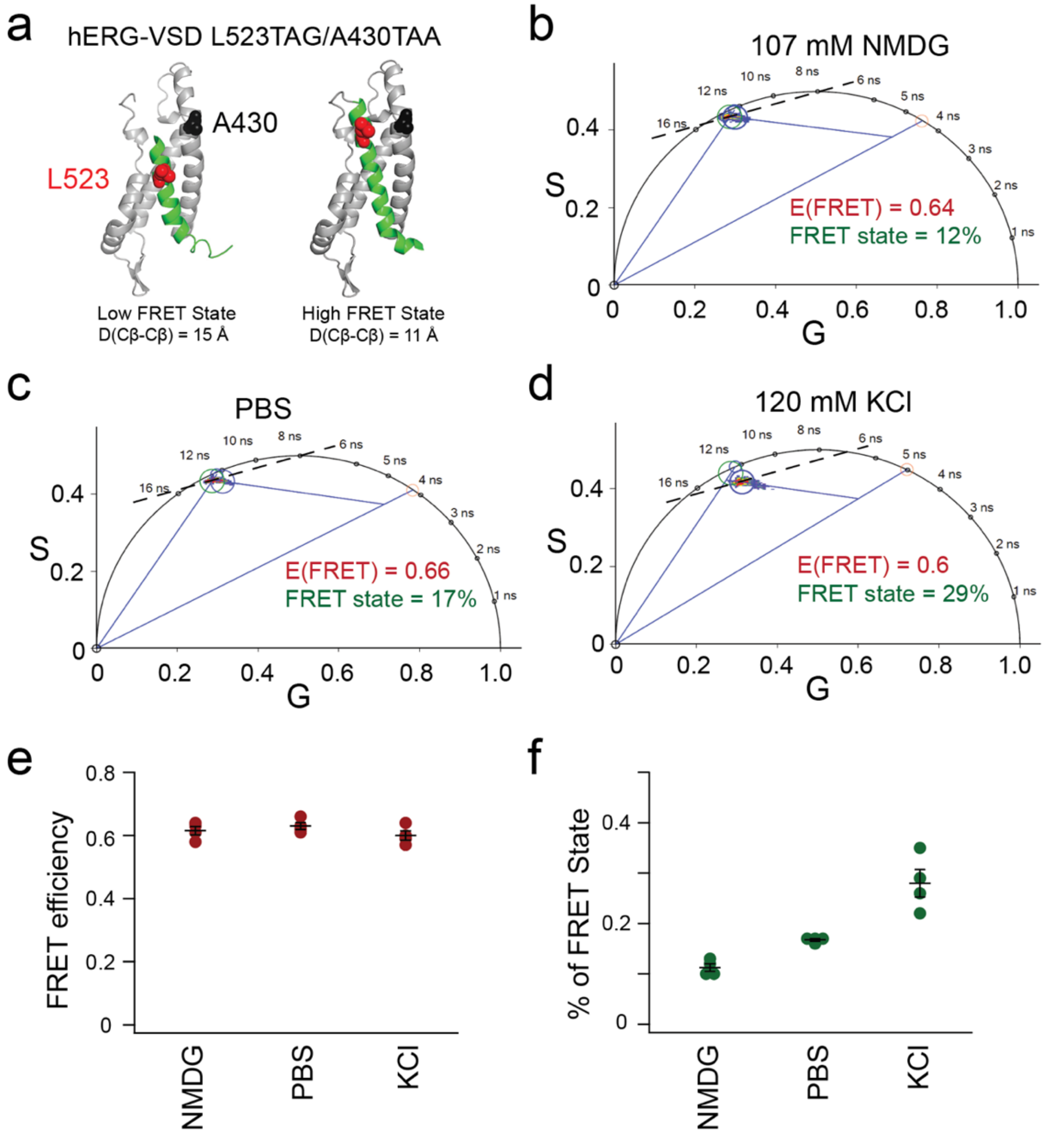


**Supplementary Fig. 6. FLIM-tmFRET on the hERG-L523TAG/A430TAA construct produced arc-shaped lifetime distribution in the phasor plot.**

**a**. Structural cartoons illustrating the low FRET state and high FRET state in the context of L523/A430 FRET pair, corresponding to the homology models of hERG-VSD in the down and up states. Different from the L523/G439 pair, the low FRET state corresponding to the S4 down conformation yields noticeable FRET.

**b-d**. Representative analysis of individual cells under various conditions indicated. The lifetime distribution has an arc shape. The primary lifetime distribution (towards shorter lifetime), after a linear fitting, yielded a notably higher FRET efficiency of approximately 0.65 (a converted distance of 10 Å) compared with that of the L523/G439 pair, indicative of a closer donor-acceptor proximity in the L523/A430 pair. In addition, the minor component of the lifetime distribution exhibited a lower FRET efficiency of 0.37, corresponding to a converted distance of 16 Å, presumably corresponding to the FRET in the down state of S4.

**e-f**. Summary data of estimated FRET efficiencies and percentage of the high FRET state. Data shown are presented as mean ± s.e.m., n = 4.

**Figure S7**

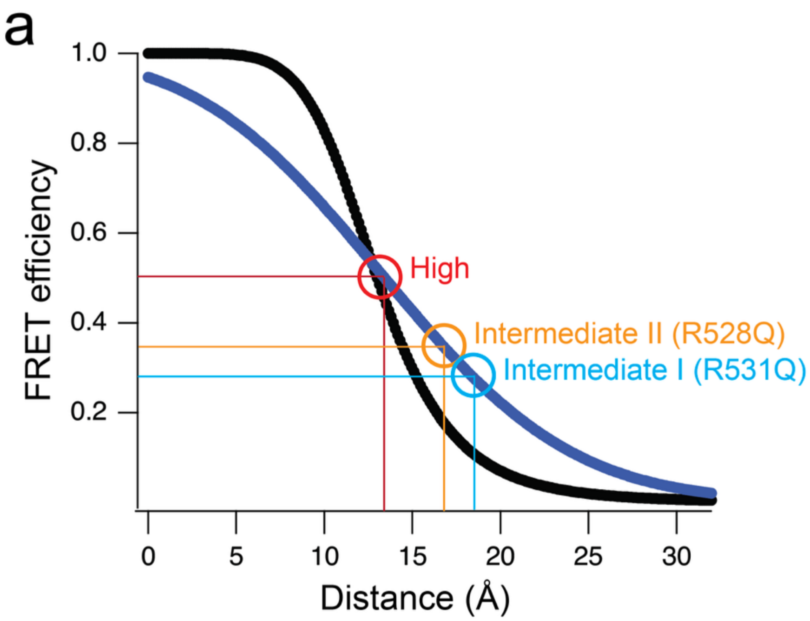


**Supplementary Fig. 7. Förster Convolved Gaussian (FCG) method of estimating distances from measured tmFRET efficiencies.**

**a.** Black curve is the standard 6$^{th}$ power dependence of FRET efficiency on donor-acceptor distance, based on Förster equation. Blue curve is the dependence based on FCG, with a standard deviation (σ) of 7.5 Å for the Gaussian distribution, taking into account of the heterogeneity of interatomic distances between FRET donors and acceptors in an ensemble FLIM measurement. The intermediate FRET efficiencies highlighted correspond to the intermediate I conformation of R531Q and the intermediate II conformation of R528Q, both are indicated as lowest-energy minima in their respective free-energy landscapes.

**Figure S8**

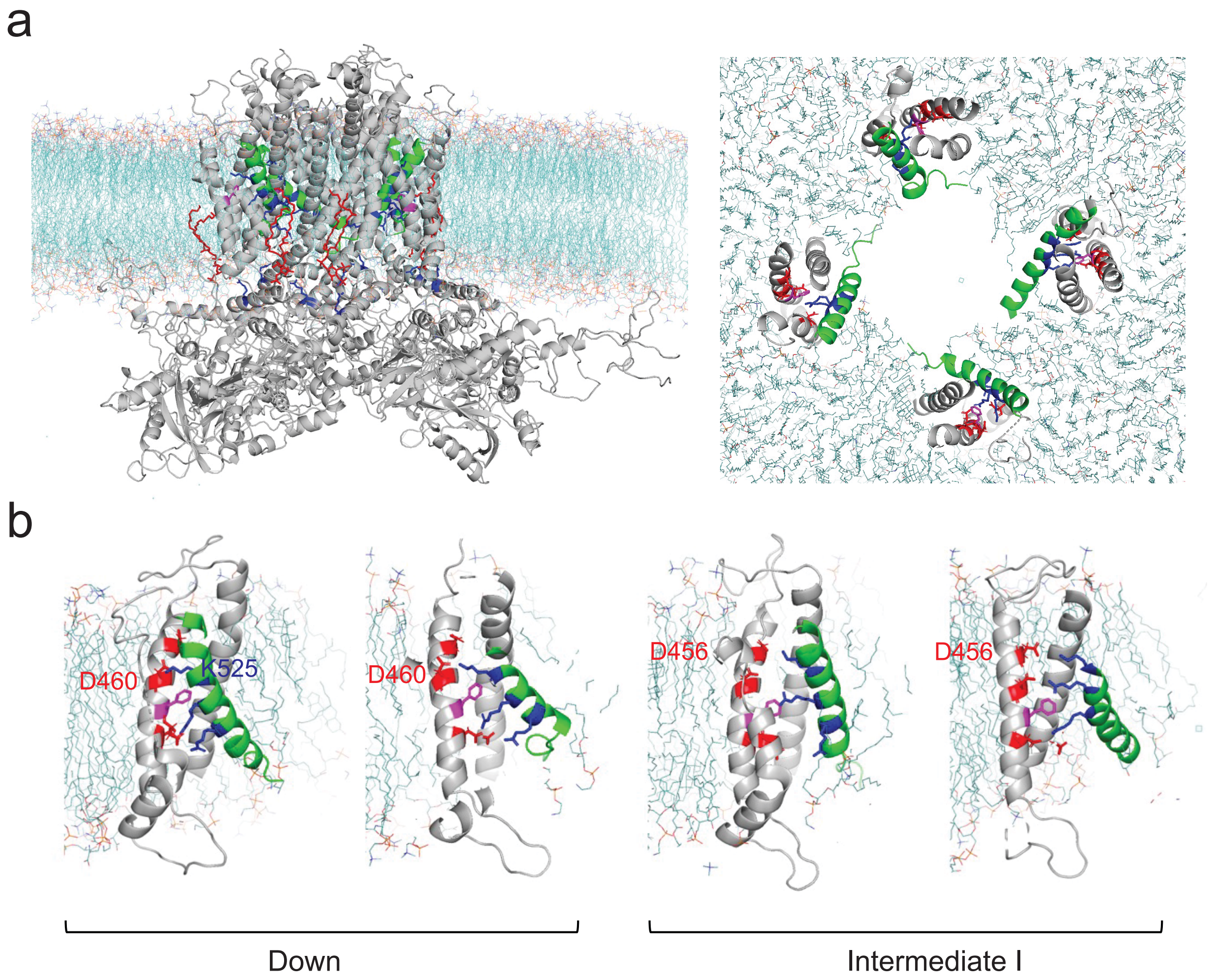


**Supplementary Fig. 8. Molecular dynamics (MD) simulation of a hERG channel tetramer in lipid membrane at hyperpolarized voltage.**

**a**. MD simulations of the S4 helix relaxing either in the down conformation or in an intermediate conformation for the four VSDs at -150 mV. PI(4,5)$P_2$ lipid (red) was included in the simulations. On the right is a top view of the four VSDs without showing the pore domain.

**b**. MD-generated structures of monomeric transmembrane domains derived from each tetrameric hERG VSD subunit, with two in the down conformation and two in the intermediate I conformation. Side views are shown in slightly different angles. In the intermediate I conformation, the K525 of S4 interacts primarily with D456, rather than with D460, in the ENC. Additionally, the D540/R665 salt bridge interaction does not appear to influence the positioning of the S4 in either the down or intermediate conformations, as a closer interaction between D540 and R665 is not specific to either conformation.

**Figure S9**

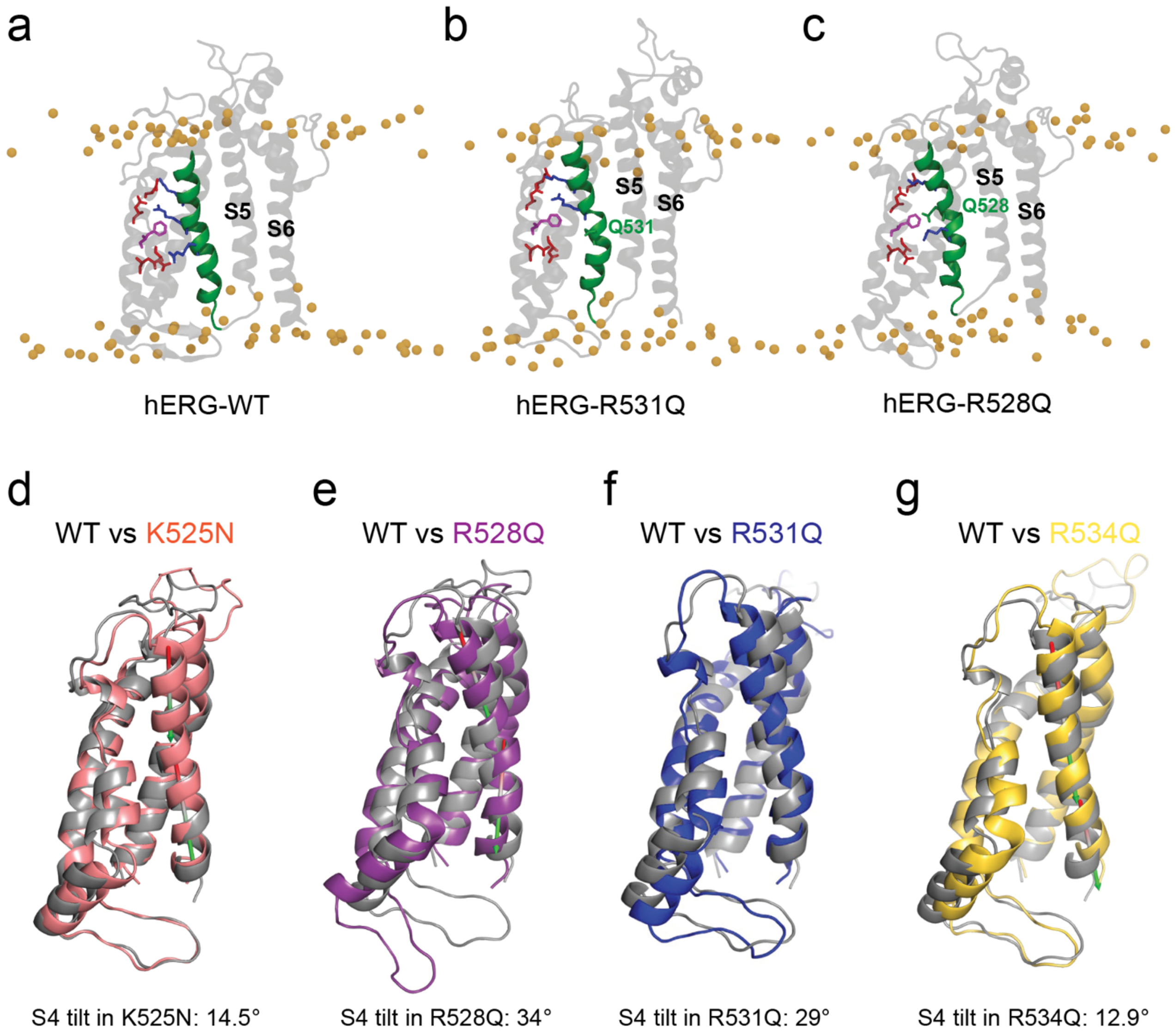


**Supplementary Fig. 9. Comparing the intermediate VSD conformations between the WT and the R531Q, R528Q mutants.**

**a-c**. MD-simulated conformations of the VSD monomer of hERG-WT (a), R531Q (b) and R528Q (c) mutants at hyperpolarized voltage (-150 mV), viewed in a lipid environment. A noticeable $3_{10}$-helical conformation, suggestive of local helix unwinding, was observed near the respective mutation sites in the S4 helix.

**d-g**. Overlay of MD-simulated structures of K525N (d), R528Q (e), R531Q (f), and R534Q (g) VSDs with the WT VSD (gray) at -150 mV. All charge-neutralizing mutations induced tilting of the S4 helix at their respective sites, with R528Q and R531Q causing markedly greater structural perturbations than K525N and R534Q. In addition, R528Q and R531Q promoted significantly more upward displacement of the S4 helix compared to the minimal shifts observed with K525N and R534Q. The tilt angles are listed for the mutant VSDs.

**Figure S10**

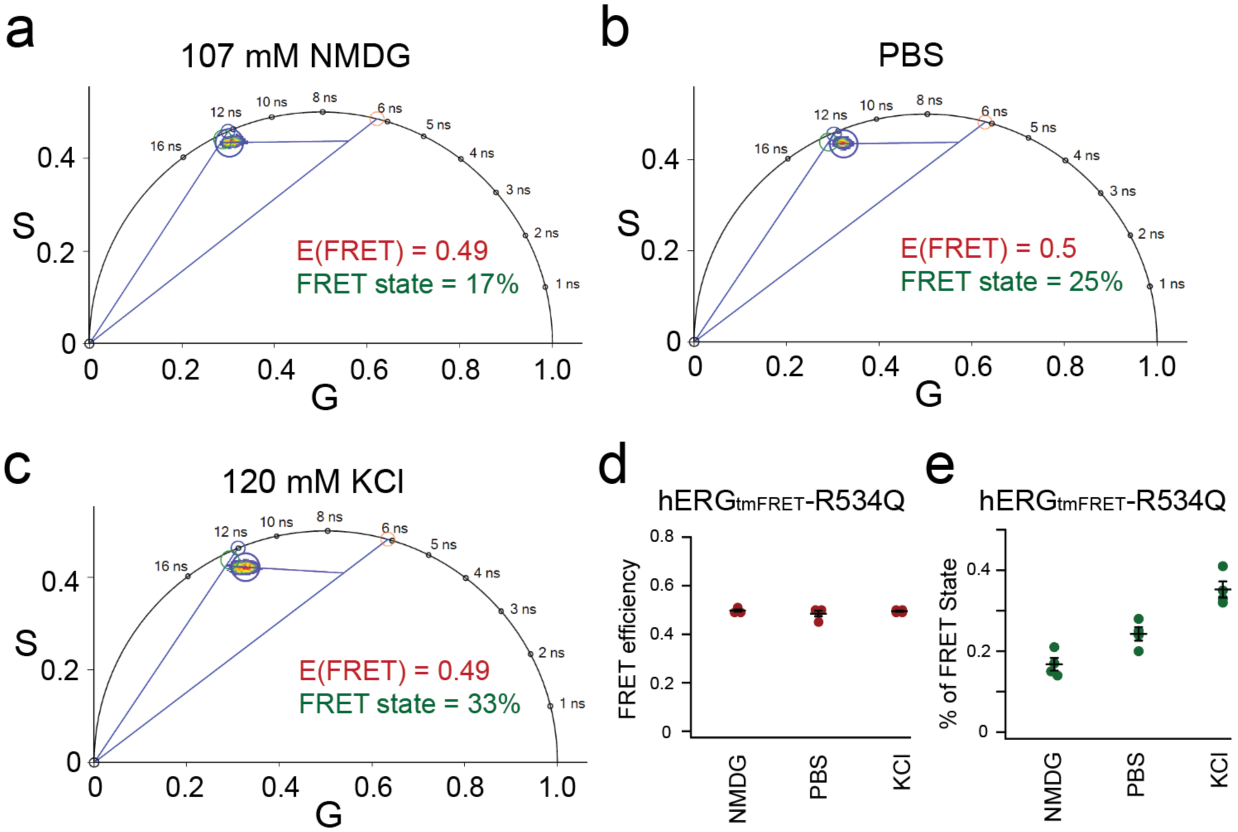


**Supplementary Fig. 10. FLIM-tmFRET on hERG$_{tmFRET}$ constructs with the incorporation of R534Q mutation.**

**a-c**. Representative analysis of individual cells under various conditions indicated. R534Q produced results similar to that in hERG$_{tmFRET}$ channels.

**d-e**. Summary data of estimated FRET efficiencies and percentage of the high FRET state using linear unmixing of the phasor approach. Data shown are presented as mean ± s.e.m., n = 4.

**Figure S11**

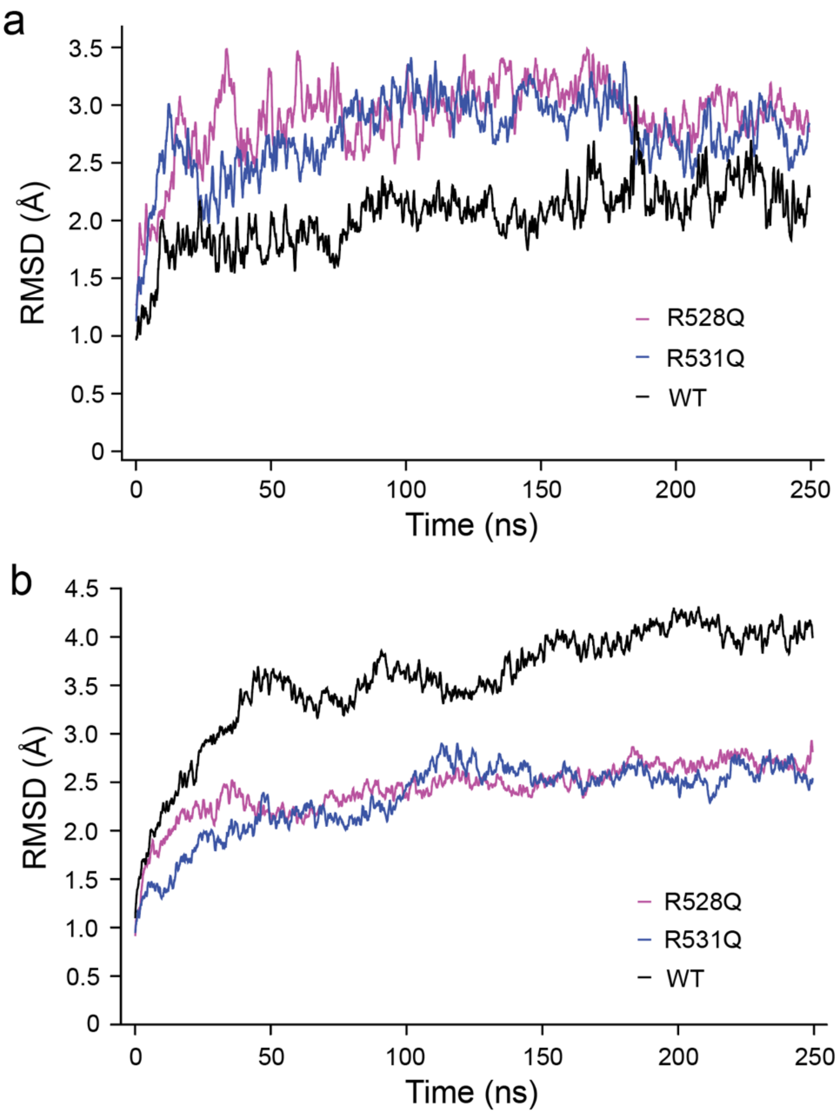


**Supplementary Figure 11. Root-mean-square deviation (RMSD) time course during equilibrium MD simulations.**

**a**. Time evolution of the RMSD for monomeric transmembrane domains of the WT, R531Q, and R528Q mutant channels, calculated relative to their respective starting conformations. These trajectories reflect the conformational stability and dynamic flexibility of individual subunits over the course of the simulation (250 ns).

**b**. RMSD profiles for the full-length tetrameric WT, R531Q, and R528Q channels, highlighting the structural stability and potential differences in global conformational dynamics within 250 ns. The mutants were generated from the equilibrated WT, therefore the change in RMSD from the start value is lower in the two mutants than in the WT.

**Table S1**. **Open probability and ΔG values for channel opening based on the probability of S4 in the activated (up) state ($P_{FRET}$), comparing the WT and the R531Q mutant.**

| Model (≥ n active) | $P_{Open}$ (WT) | $P_{Open}$ (R531Q) | ΔG (WT) (kcal/mol) | ΔG (R531Q) (kcal/mol) |
|---|---|---|---|---|
| $P_{FRET}$ | 0.35 | 0.20 | 0.35 | 0.20 |
| ≥1 subunit | 0.821 | 0.590 | 0.37 | 0.82 |
| ≥2 subunits | 0.437 | 0.181 | 0.73 | 1.64 |
| ≥3 subunits | 0.126 | 0.027 | 1.10 | 2.46 |
| ≥4 subunits | 0.015 | 0.002 | 1.47 | 3.28 |
